# Phenome-wide genetic framework to identify mechanisms of social effects

**DOI:** 10.64898/2026.03.10.710784

**Authors:** Hélène Tonnelé, Francesco Paolo Casale, Amelie Baud

**Author notes:** Corresponding author: Amelie Baud.

## Abstract

Phenotypes are shaped not only by an individual’s genotype (direct genetic effects, DGE) and environment but also by the genetic composition of social partners through indirect genetic effects (IGE). Although IGE have been detected across many traits and species, their mechanisms remain largely unknown, particularly for physiological traits. Here we introduce a phenome-wide genetic framework that identifies proxy phenotypes for the heritable traits of social partners mediating IGE by estimating genetic correlations between IGE on focal phenotypes and DGE on measured traits. Applying this approach to two large, outbred mouse datasets comprising hundreds of behavioural and physiological phenotypes, we find that behavioural traits are neither more affected by IGE nor better proxies for the traits mediating them, challenging the prevailing behavioural-centric view of social effects. Instead, immune, metabolic and growth phenotypes are both affected by IGE and informative proxies for their underlying mechanisms, potentially reflecting the social transmission of gut microbes. Our framework provides a novel strategy to better understand the genetic basis of complex traits and uncover mechanisms of social effects.

## Introduction

Phenotypes do not arise in isolation but are shaped by the social environments in which individuals develop and live. Across mammals, interactions with conspecifics influence a wide range of biomedical traits, from early development to adult behaviour and physiology^1^. Yet, identifying the mechanisms underlying social effects remains challenging, for several reasons: first, it is difficult to know *a priori* all the traits of social partners that could influence the phenotype of interest. Secondly, socially interacting individuals tend to share resources and exposures, leading to pervasive correlations between their phenotypes^2^. Disentangling genuine social influences from confounding effects therefore requires approaches that can move beyond descriptive correlations.

Indirect genetic effects^3,4^ (IGE) are social effects of genetic origin. They arise when heritable traits of social partners influence the phenotype of interest (Fig. 1A). There are several advantages to using IGE to study social effects. First, IGE can be detected without specifying the traits of social partners involved, by testing for associations between the phenotype of interest and the genotypes of social partners^5^. Secondly, IGE can serve as causal anchors to establish causal relationships since the genotypes of the social partners are assigned at birth and unaffected by environmental factors and phenotypes^6^. Finally, modelling IGE is essential to understand the evolution of socially affected phenotypes^3,7^.

**Figure 1.**
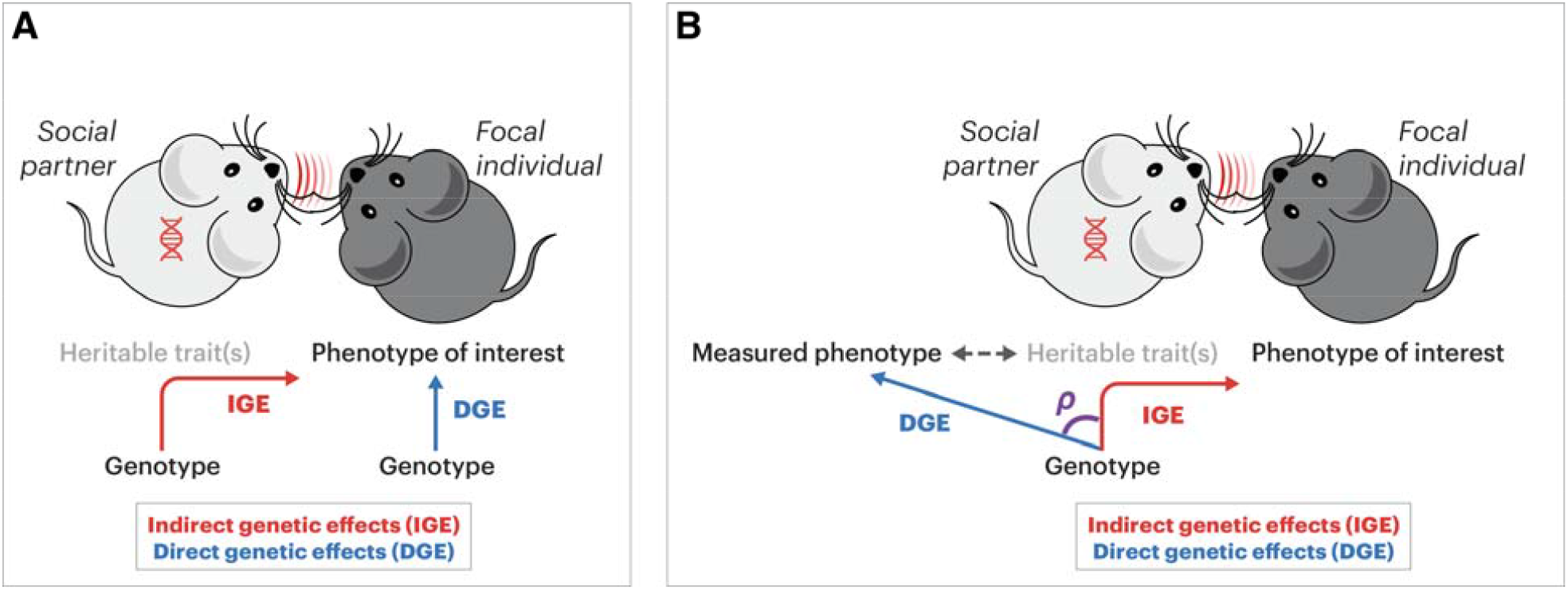
Quantitative genetics framework to uncover new mechanisms of indirect genetic effects. (A) Definition of key terms. The heritable trait(s) of social partners mediating IGE are unobserved. (B) Representation of the genetic correlation ρ between IGE on the phenotype of interest and DGE on a measured phenotype to identify good proxy phenotypes for the heritable traits of social partners mediating IGE.

IGE have been uncovered across a variety of animal species, types of relationships (e.g. between parents and offspring, between males and females, between peers, etc.) and phenotypes^4,8,9^. Behavioural phenotypes are most commonly reported to be affected by IGE^9,10^. Strong IGE are easy to understand for social behaviours (i.e. behaviours that are only expressed when social partners are present) and behaviours that are “contagious”, as individuals genetically predisposed to a high trait value can elicit higher trait values in their partners. For example, *Peromyscus* mice genetically predisposed to aggression can instigate higher aggression in others^11^ and an individual’s smoking behaviour is influenced by peer smoking^12^.

Beyond behavioural traits, IGEs have been reported on a wide range of developmental, physiological, morphological, reproduction and survival phenotypes^9^. In humans, robust IGE in eighty thousand couples of the UK Biobank were reported for educational attainment and arthrosis^13^. In laboratory mice, IGE were shown to affect biomedical markers, including immune, metabolic and growth phenotypes^14,15^. Although IGE are an important source of phenotypic variation, the underlying mechanisms remain unknown, particularly where non-behavioural phenotypes are concerned.

To address this gap and leverage the increasingly large number of biobanks available for genetics research^16–20^, we introduce a genetic framework that identifies good proxy phenotypes for the traits of social partners influencing a phenotype of interest on the basis of their genetic correlation ρ with the traits mediating IGE (Fig. 1B). Genetic correlations can arise through causal relationships or pleiotropy and are widely used to identify functional relationships between phenotypes^21^. We apply this strategy to two datasets collected in thousands of “HS” and “CFW” outbred laboratory mice and including a wide range of behavioural, physiological, and morphological phenotypes relevant to human diseases^22–24^ (Fig. 2 and Supplementary Tables 1 and 2). In these datasets, we previously detected IGE arising from the genotypes of cage mates on a wide range of phenotypes, many of which were not previously known to be affected by social effects^14,15^. In most cases, the mechanisms of social effects are completely unknown. A better understanding of the traits of social partners mediating social effects could open the door to preventive or therapeutic interventions based on a better management of social interactions^25^.

**Figure 2.**
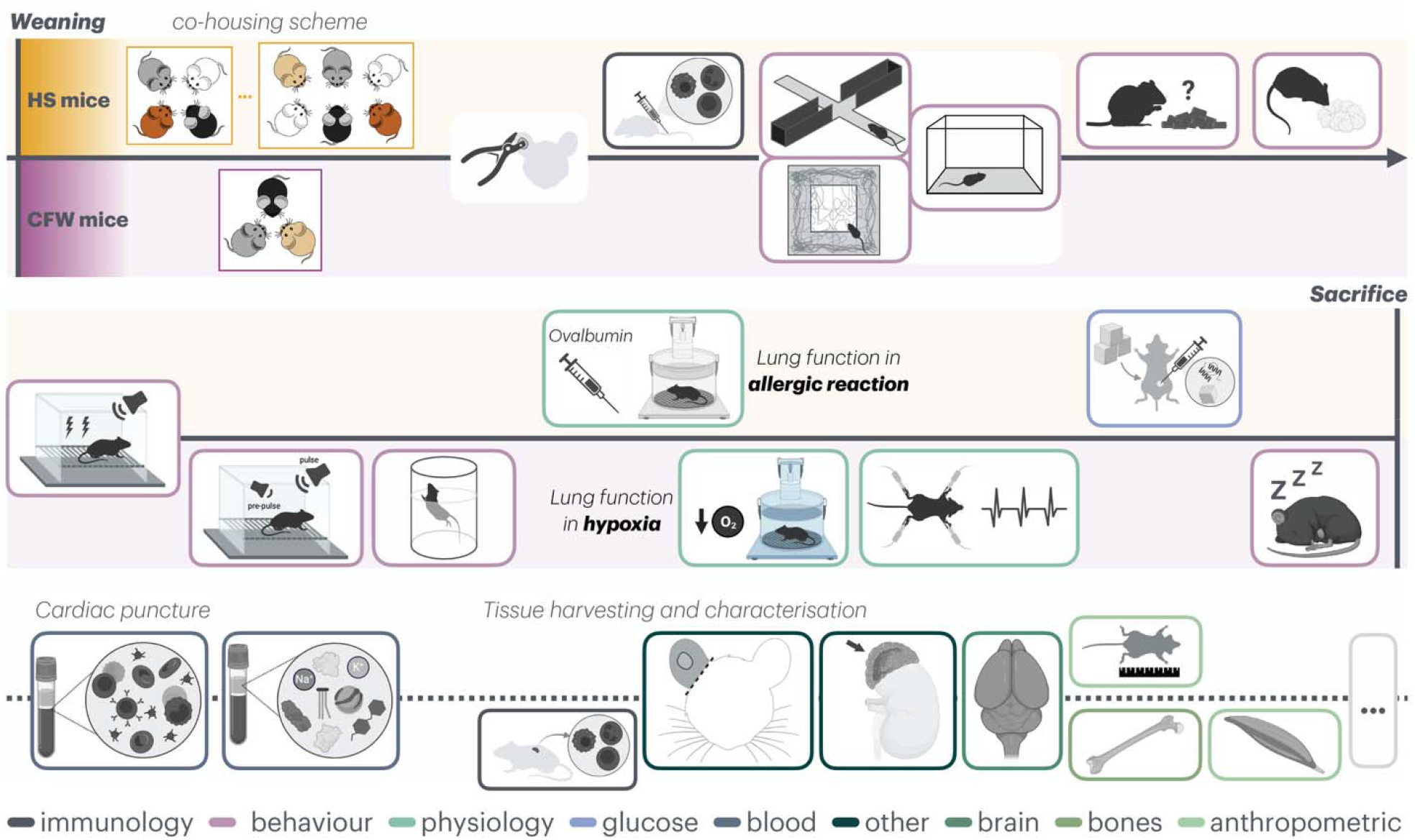
Phenome-wide data collected in HS and CFW mice. Tests performed in both populations are displayed along the grey timeline, while those specific to a single population are shown above (HS mice) or below (CFW mice) the line. The colour of the border line of each box indicates the category of the phenotypes collected (see also Supplementary Tables 1 and 2).

## Results

### Refining the leading view that behaviours are more affected by IGE than other phenotypes

Our previous quantification of IGE in the HS and CFW mouse datasets suggested that behaviours were not the class of phenotypes most strongly affected by IGE^14,15^, contrary to the prevailing view in the field and the results of a recent meta-analysis that did not include the HS and CFW datasets^9^. To address this apparent discrepancy, we systematically compared IGE on behavioural and non-behavioural phenotypes in the mouse datasets and re-analysed the data of the meta-analysis to distinguish between social and non-social behaviours, because the mouse datasets did not include social behaviours. In mice, we found no evidence that behaviours were more affected by IGE than non-behavioural phenotypes (one-tailed non-parametric Wilcoxon rank-sum test *p*-value = 0.91 for HS mice, *p*-value = 0.94 for CFW mice, Fig. 3A and 3B). In the meta-analysis, we found that the large IGE observed for behaviours were driven by social behaviours (Fig. 3C), resolving the apparent discrepancy.

**Figure 3.**
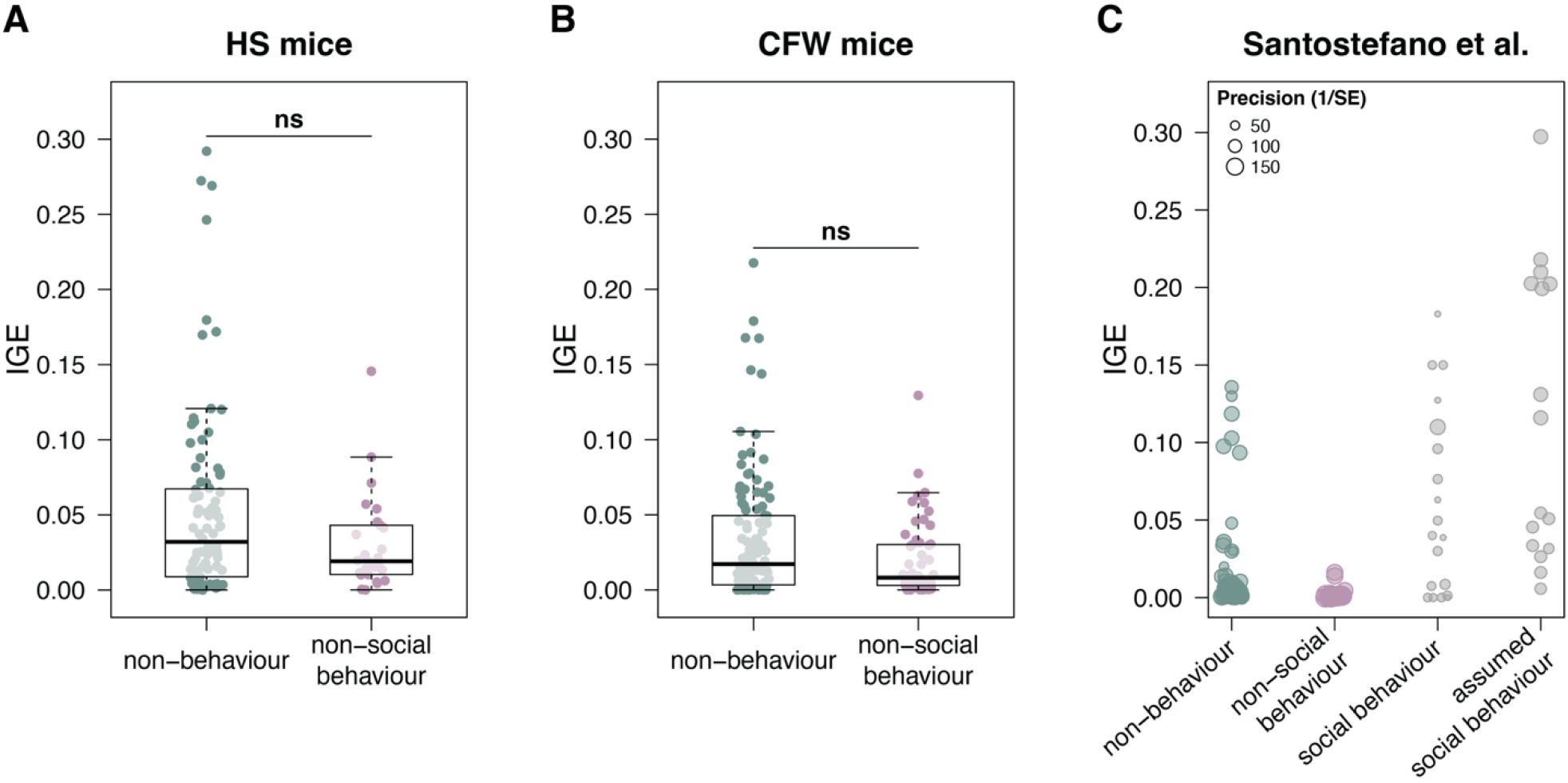
Comparison of the magnitude of indirect genetic effect (IGE) between behavioural and non-behavioural phenotypes. (A) Our estimates from HS outbred laboratory mice. (B) Our estimates from CFW mice. Each dot represents an individual trait, coloured in green if non-behavioural or in pink if behavioural. (C) IGE estimates from Santostefano et al.^9^, shown for non-behavioural, non-social behavioural, social behavioural, and assumed social behavioural traits; point size reflects estimate precision (inverse standard error). Dots are coloured grey for phenotypic classes that are not present in our study.

### Validation of a new implementation of bivariate IGE models

To identify good proxies for the traits of cage mates mediating IGE, we implemented a bivariate linear mixed model that jointly models direct and indirect genetic effects across pairs of phenotypes. This model allows us to estimate the genetic correlation ρ between IGE on the phenotype of interest and DGE on each measured phenotype. A high correlation suggests that the measured phenotype is genetically related to the unobserved traits of cage mates mediating IGE (Fig. 1B). We used simulations based on the real genotypes and cage structure of the HS and CFW datasets to confirm that the bivariate model yields unbiased genetic estimates and well-calibrated *p*-values (Supplementary Figs. 1 and 2).

### Behaviours are not better proxies for the traits mediating IGE than non-behavioural phenotypes

Because social partners are often thought to exert their influence through behaviours, we first tested whether the behavioural phenotypes included in the HS and CFW datasets are better proxies for the traits mediating IGE than the other, non-behavioural phenotypes. We limited our analysis to the phenotype pairs where IGE on one phenotype and DGE on the other phenotype were strong enough (>5% of phenotypic variation explained). We saw no evidence of this, as the correlations ρ involving DGE on behavioural phenotypes were no greater than the correlations involving DGE on non-behavioural phenotypes (one-tailed non-parametric Wilcoxon rank-sum test *p*-value = 0.33 and 1.0 for behavioural and non-behavioural phenotypes, respectively, in HS mice; *p*-value = 0.57 and 0.85 for behavioural and non-behavioural phenotypes, respectively, in CFW mice; Fig. 4). Thus, behaviours are no better proxies for the traits mediating IGE than non-behavioural phenotypes.

**Figure 4.**
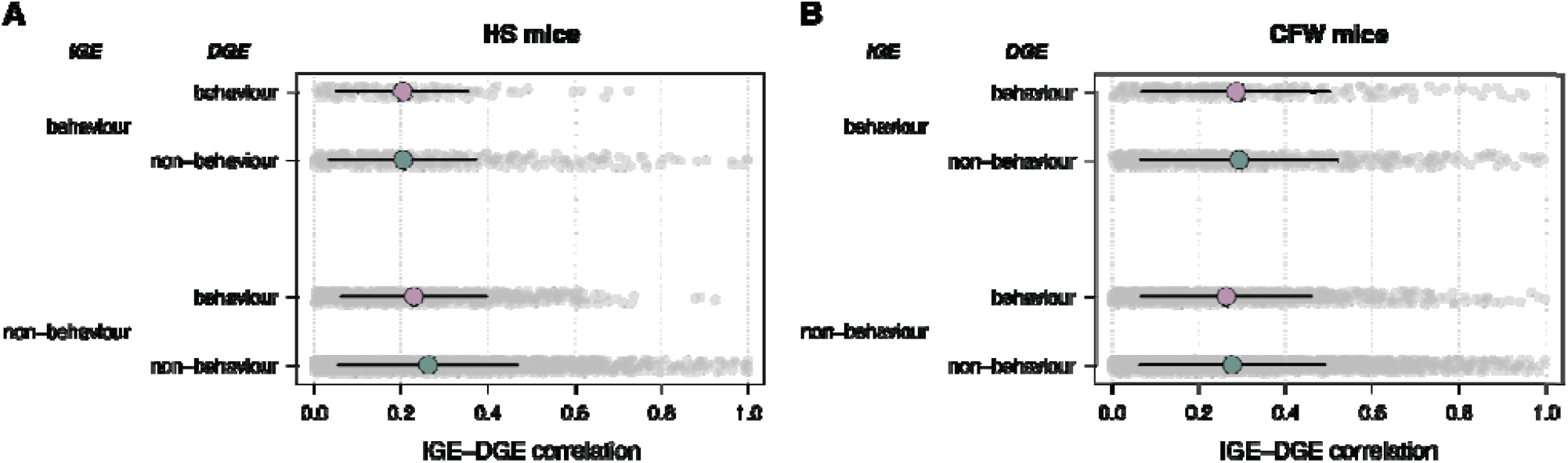
Behavioural VS non-behavioural phenotypes as proxies for the traits of cage mates mediating IGE. The comparison was done in both HS mice (A) and CFW mice (B) and separately for IGE on behavioural phenotypes (top two rows) and IGE on non-behavioural phenotypes (bottom two rows). Each dot represents the IGE-DGE correlation ρ for a pair of phenotypes; the coloured dots and black lines show the mean and standard deviations for each distribution.

### Identifying good proxies for the heritable traits of cage mates mediating IGE

To gain insights into the traits of social partners mediating IGE, we visualized the IGE-DGE correlations ρ in a heat map where phenotypes of interest (rows) and potential proxy phenotypes (columns) were clustered based on the significance of the IGE-DGE correlation (ρ ≠ 0). We limited our analyses to the phenotype pairs where IGE on one phenotype and DGE on the other phenotype were strong enough (>5% of phenotypic variation explained). In HS mice (Fig. 5A), we observed a cluster featuring immune, metabolic and growth phenotypes as both traits affected by IGE and good proxies for the traits mediating these IGE. More specifically, the immune phenotypes included counts and size of T and B lymphocytes, expression of CD4 at the surface of T cells, and measures of airway resistance following immunization with ovalbumin. Metabolic traits included glycemia before and 15 minutes after intraperitoneal glucose injection, and serum ion levels. The growth phenotypes were measures of body weight collected throughout the life of the mice, as well as adult body length. Because many of the immune and metabolic phenotypes were blood and serum phenotypes, we wondered whether the clustering observed could be due to batch effects associated with the blood draws and subsequent processing. Since the blood and serum phenotypes were derived from four separate blood draws spread across the life of the mice and four separate analytical procedures (see Methods), we considered that batch effects were extremely unlikely to explain the clustering and instead concluded that the high genetic correlations observed arose from causal relationships or pleiotropy, but not experimental confounding.

**Figure 5.**
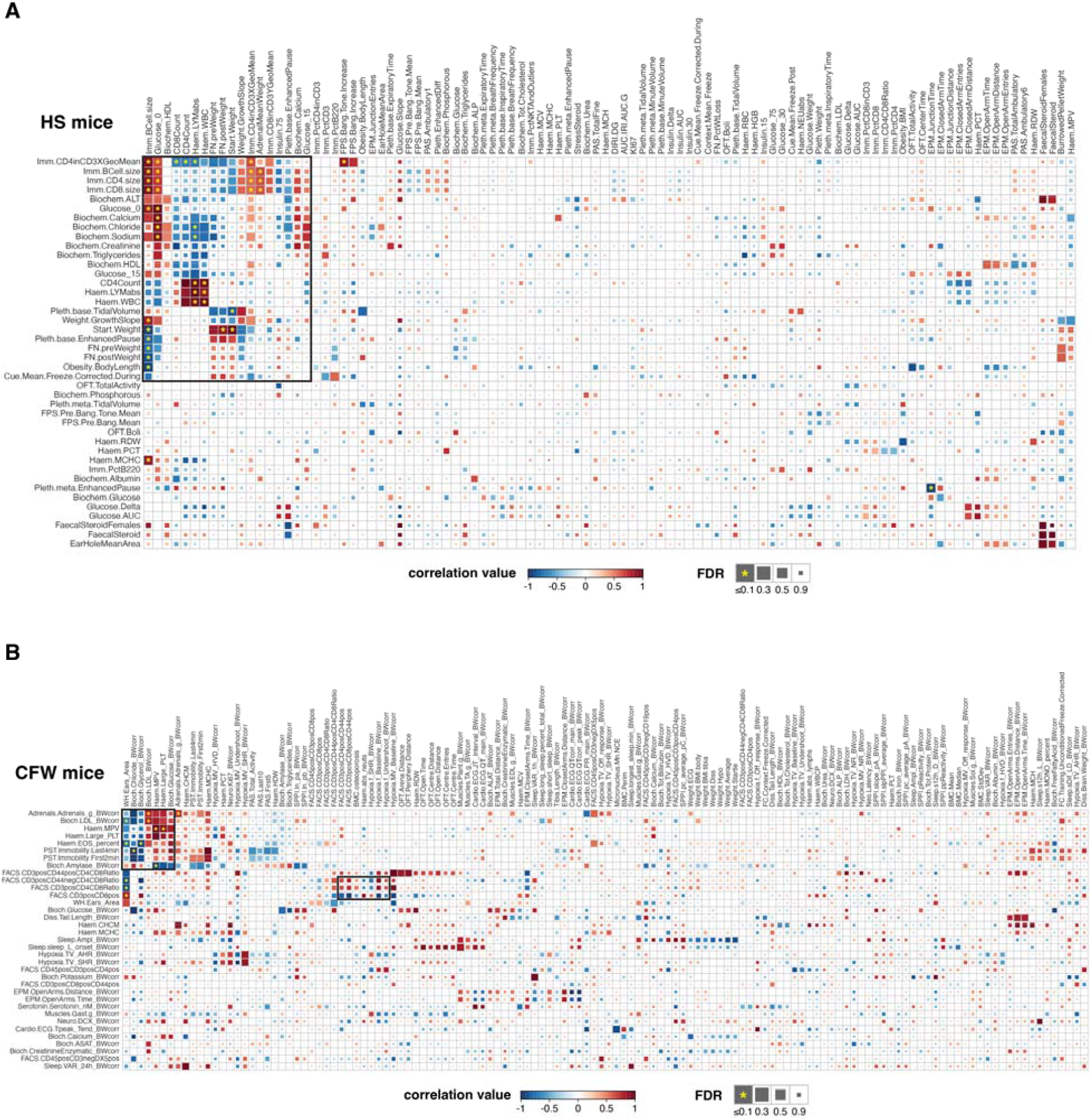
Clustered heat maps of IGE-DGE correlations ρ in (A) HS and (B) CFW mice. Phenotypes affected by IGE (rows, >5% of phenotypic variation explained) and potential proxy phenotypes affected by DGE (columns, >5% of phenotypic variation explained) were clustered based on the significance of the IGE-DGE correlation (ρ ≠ 0), which was calculated using a likelihood ratio test (see Methods). In HS mice (A) the black square highlights a cluster of immune, metabolic and growth phenotypes; in CFW mice (B) the black square on the top left highlights a large, heterogeneous cluster, whereas the square in the middle highlights a smaller cluster of T lymphocytes phenotypes.

In CFW mice, IGE-DGE correlations were generally less significant, but we nevertheless observed two clusters (Fig. 5B). The larger one was highly heterogeneous in terms of the types of the phenotypes involved. The smaller one featured T lymphocytes counts and ratios as phenotypes affected by IGE and proxy phenotypes for the traits mediating these IGE (Fig. 5B). Thus, the smaller cluster is reminiscent of the cluster observed in HS mice (Fig. 5A).

Altogether, these results indicate a shared genetic architecture between DGE and IGE affecting T lymphocyte phenotypes. In HS mice, these genetic effects also influence metabolic and growth phenotypes directly and indirectly (i.e. through social effects).

### Similar traits mediate IGE in groups of males and groups of females

In our analyses we implicitly modelled IGE as arising from the same traits of cage mates in groups of males and groups of females. To evaluate this assumption, focusing on a single phenotype of interest, we split the phenotype between a male phenotype and a female phenotype, assigning missing values to females for the male phenotype and *vice versa*. Doing so, the genetic correlation between IGE on the male phenotype and IGE on the female phenotype reflects the extent to which similar genetic variants, hence similar biological pathways, give rise to the traits of social partners mediating IGE in groups of males and in groups of females. After confirming using simulations that genetic estimates and *p*-values were also valid in this analysis (Supplementary Fig. 3 and Supplementary Fig. 4), we estimated the male-female genetic correlation for each phenotype included in the dataset and tested whether it was different from one, which would provide evidence that different traits of social partners underlie IGE in groups of males and groups of females. The correlations ranged from 0.29±0.56 to 0.99±0.36 in HS mice (mean 0.87) and from -0.27±0.43 to 1±1.21 in CFW mice (mean 0.63). No correlation was significantly different from 1 (FDR > 0.1), providing no evidence that different traits of social partners mediate IGE in male and female groups.

## Discussion

In this study, we introduce a phenome-wide, genetics-based strategy to gain insights into the traits of social partners mediating indirect genetic effects (IGE). By leveraging genetic correlations between IGE on phenotypes of interest and direct genetic effects (DGE) on a broad array of measured traits, we identify phenotypes that act as informative proxies for the heritable traits of social partners influencing these phenotypes of interest. Applying this approach to two independent outbred mouse populations provided evidence that similar genetic effects act on adaptive immunity phenotypes directly and indirectly. This signal was visible in both datasets, and in HS mice these genetic effects were also correlated with DGE and IGE on metabolic and growth phenotypes.

### Revisiting the behavioural-centric view of social genetic effects

IGE are most intuitively understood for social behaviours (i.e. behaviours that are expressed only in the presence of other individuals) and behaviours that are contagious, because individuals genetically predisposed to a high trait value can elicit higher trait values in their partners (positive feedback loops). Consistent with this intuition, behavioural traits have dominated the IGE literature and were concluded to be particularly affected by IGE in a recent meta-analysis. However, by re-analysing the data from the meta-analysis with a finer distinction between social and non-social behaviours, and by analysing two independent mouse datasets that included contagious behaviours (anxiety and depressed mood/stress coping strategy^26^) but no social behaviours, we showed that the tendency for behaviours to be strongly affected by IGE is driven by social behaviours. Our findings therefore refine our understanding of IGE on behavioural phenotypes. Furthermore, in the mouse datasets non-behavioural phenotypes exhibited IGE of comparable—or even greater—magnitude as non-social behavioural phenotypes, and the best proxies for traits of social partners mediating IGE were non-behavioural traits. These results call for a broader conceptualization of social genetic effects, in which behaviours represent only one of several possible conduits through which genetic variation in social partners can influence focal individuals. Importantly, this broader view is necessary to explain the strong and reproducible IGE observed for physiological phenotypes, such as immune traits, metabolism, and growth phenotypes.

### Phenome-wide genetic correlations as a window into social mechanisms

The central methodological advance of this work is the use of phenome-wide genetic correlations between IGE and DGE to identify measured traits that are good proxies for the unobserved traits mediating social effects. This strategy builds on the logic of genetic correlation analyses widely used to study shared disease aetiology^21^, which exploits the causal anchoring provided by genotypes to reduce confounding relative to phenotypic correlations. Although genetic correlations do not, by themselves, distinguish between causality and pleiotropy, both mechanisms are biologically meaningful in the context of IGE, as they reflect shared heritable pathways that shape social environments and influence evolutionary dynamics. The validation of our bivariate IGE models through extensive simulations confirms that these correlations can be estimated with minimal bias, even in complex social designs. This opens the door to systematic, unbiased scans of phenome datasets to generate mechanistic hypotheses about social effects, without requiring prior knowledge of the relevant traits or behaviours. Such phenome datasets are increasingly available in humans, laboratory, wild and agricultural populations.

### Gut microbiota transmission as a plausible mechanism for physiological IGE

Across both mouse populations, behavioural phenotypes were not better proxies for the traits mediating IGE than non-behavioural phenotypes. Instead, the most robust signal emerged from T lymphocytes phenotypes, which were affected by IGE and good proxies for the traits of social partners mediating these IGE in both datasets. In HS mice, DGE and IGE on metabolic and growth phenotypes also shared a genetic basis with DGE and IGE on immune phenotypes. Blood lymphocytes are very unlikely to be passed between mice sharing a cage, and growth phenotypes simply cannot. Hence, these phenotypes are merely proxies for the traits mediating IGE: they are related to those traits through causal relationships or pleiotropy. Which factors, then, can be passed between mice sharing a cage and affect blood lymphocytes levels and, also, metabolic and growth phenotypes? A plausible hypothesis is commensal gut bacteria, which are well-known to be transferred between mice sharing a cage through allo-coprophagy and affect the immunity, metabolism, and growth of their host (Fig. 6). This hypothesis is further supported by recent evidence from our lab that demonstrated IGE on a subset of gut bacteria in laboratory rats^27^. Alternatively, stress and its transmission could explain these signals. The lack of correlation between DGE on the anxiety and stress-coping phenotypes included in both mouse datasets^28^ and IGE on immune, metabolic and growth phenotypes, however, does not support this alternative hypothesis. Future work integrating experimental manipulations of the traits suspected to mediate IGE—such as targeted manipulations of the gut microbiota of one mouse and evaluation of the immune, metabolic and growth phenotypes of the other mice in the cage—will be essential to directly test the proposed mechanisms.

**Figure 6.**
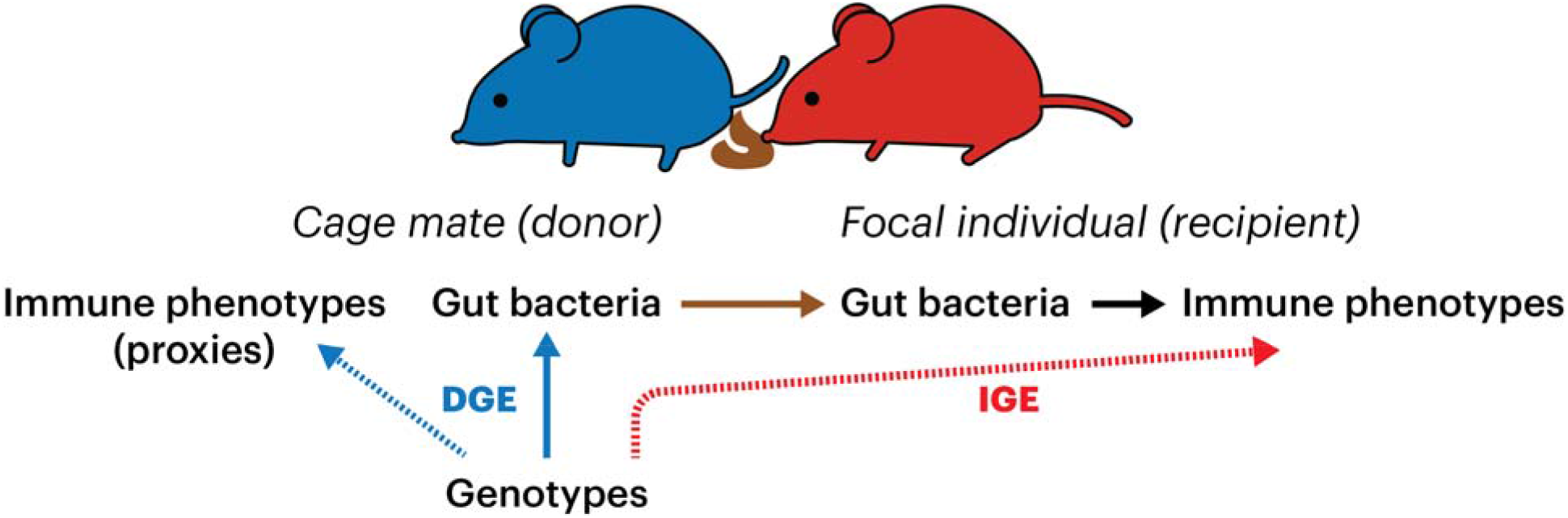
Hypothesized mechanism for the high correlations between IGE and DGE on immune phenotypes (dashed lines): gut microbiome transfers through allo-coprophagy.

Many animals besides rodents are coprophagic and ingest faeces from conspecifics^29^. Humans are not coprophagic, yet there is ever increasing evidence of pervasive horizontal transmission of commensal gut microbes in our species, including within the household and at nurseries^30–33^. The social transmission of gut microbes could mediate social effects, including IGE, on immune, metabolic and growth phenotypes in other animal species, including ours.

### Similar social mechanisms in males and females

We formally compared IGE in male-only and female-only groups and found no evidence that different mechanisms are at play in the two sexes, justifying the analysis of IGE in males and females jointly. There are well-established differences in social behaviours between males and females in mice, including in the laboratory setting, but since (social) behaviours do not seem to be the main mechanisms for IGE in our study, the lack of sex differences in the mechanisms of IGE is not unexpected.

### Limitations and future directions

Several limitations of this study warrant consideration. First, the phenotypes available in HS and CFW mice do not include measures of social behaviours such as aggression or affiliation, limiting our ability to implicate social behaviours in IGE. Secondly, albeit similar, the phenotypes included in the HS and CFW mouse datasets were not the same, limiting our ability to identify mechanisms that generalise across genetic backgrounds and experimental designs, hence are more robust. Finally, in the HS dataset only, DGE and IGE were confounded with maternal (genetic) effects. We checked that the IGE-DGE correlations analysed in this study were largely unaffected by this confounding (Supplementary Fig. 5).

Despite these limitations, our results demonstrate the ability of phenome-wide, genetics-based approaches to fundamentally expand the exploration of social effects beyond behaviour. As increasingly rich phenotypic and multi-omics datasets become available in both animal models and humans, this strategy will become widely applicable, with potential implications for medicine, conservation, and animal breeding.

## Methods

### Statistical models with direct and indirect genetic effects

#### *Univariate models* (used for Fig. 3)

The following model, which is the same as the models we used in two of our previous studies of IGE^41,42^, was used to quantify DGE, IGE and their correlation, while accounting for non-genetic factors contributing to phenotypic variation:

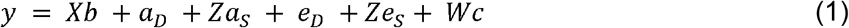

*y* is the vector of phenotypic residuals (phenotype of interest), *X* is a vector describing how many rats are in each cage and *b* the corresponding estimated fixed effect, *W* is the matrix of cage assignments and *c* the corresponding vector of random cage effects. *a*_*D*_ is the vector of random additive DGE, *a*_S_ is the vector of random additive IGE and *Z* is the matrix indicating, for each mouse, which are the cage mates (importantly *Z*_*i,i*_ = 0). *e*_*D*_ and *e*_*s*_, also random effects, refer to the non-genetic component of direct and indirect effects.

The joint distribution of all random effects was defined as:

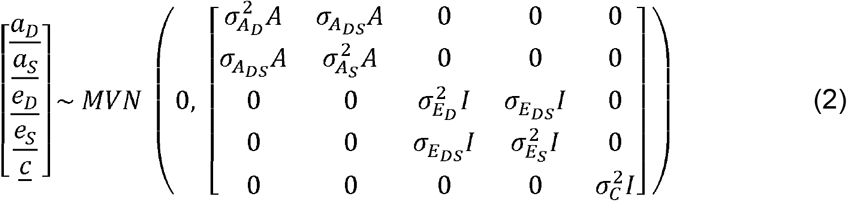

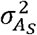 reflects the proportion of phenotypic variance explained by IGE and is the estimate reported in Fig. 3.

#### *Bivariate models* (used for Figs. 4 and 5)

The main estimate of interest in this study is the genetic correlation ρ between IGE on a phenotype of interest and DGE on a measured phenotype that is being evaluated as a potential proxy phenotype. The joint distribution of all random effects in the bivariate model was defined as:

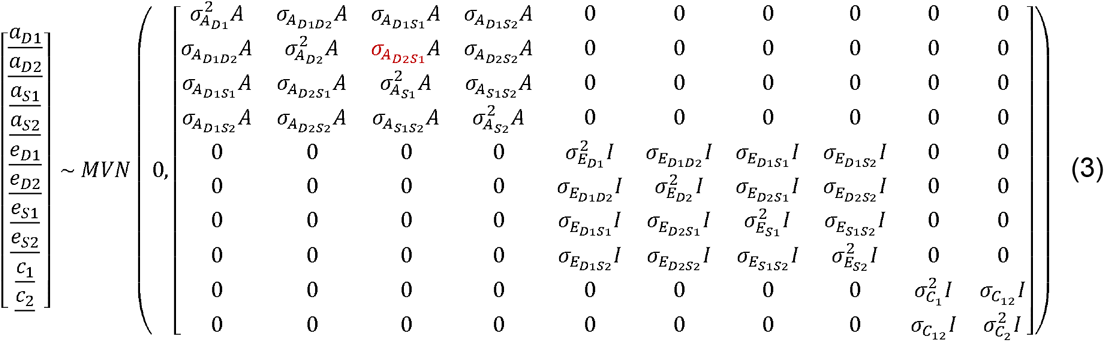

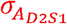 corresponds to the correlation of interest, which is reported in Fig. 4 and Fig. 5 and called 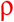 throughout this manuscript.

To evaluate whether IGE in groups of males and groups of females arise from different mechanisms, we used the same bivariate model but, in that case, one of the outcomes (*y*_1_) was the female phenotype (males had missing values) and the other (*y*_2_)) the male phenotype (females had missing values). For this analysis only, 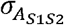 was the estimate of interest.

The model was developed within a flexible linear mixed-model framework^34,35^ and parameter estimation was performed by restricted maximum likelihood using gradient-based optimisation.

Significance values for the correlations were obtained using likelihood ratio tests, comparing a model with unconstrained correlations to a model in which the correlation of interest was fixed to the value tested (0 for 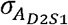 and 1 for 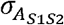). The LRT statistic for each test was compared to the chi-square distribution with one degree of freedom to get the *p*-value. To account for multiple testing, we controlled the false discovery rate using the function p.adjust from R stats package.

### Description of the mouse datasets

Heterogeneous Stock (**HS**) mice: we used the same cages, phenotypes, and genetic relatedness matrix as in Baud et al.^14^. The data were originally published by Valdar et al.^22,23^ (genotypes and organismal phenotypes) and Huang et al.^36^ (organismal phenotype: cellular proliferation in the subgranular zone of the dentate gyrus, a measure of adult neurogenesis). In brief, 2,448 male and female mice were included in this study. They were related at various levels, housed in same-sex groups of 2 to 7 mice (but mostly 3 and 4 mice), and the groups included many (but not only) full siblings. Genotypes at 13,459 single nucleotide polymorphisms were available for a subset of 1,940 mice, and pedigree data for all mice. The genetic relatedness matrix is the single-step (or H) matrix, constructed using both the genotypes and the pedigree. Some phenotype data were available for all 2,448 mice, but the number of mice phenotyped for each organismal phenotype varied. Covariates such as sex, body weight, and group size were considered and phenotypes normalised using a covariate-aware Box-Cox transformation. The covariates were fitted as fixed effects in all analyses of HS data.

Swiss Webster (Crl:CFW(SW)-US_P08, **CFW**) mice: we used the same cages, phenotypes and genetic relatedness matrix as in Baud et al.^15^. The data were originally published by Nicod et al.^24^. In brief, 1,812 male and female mice were included in this study. They were unrelated to one another (beyond the baseline relatedness associated with the maintenance of the colony) and housed in same-sex groups of 3 mice. Genotypes at 353,697 LD-pruned single nucleotide polymorphisms were available for all mice and used to build the genetic relatedness matrix. Some phenotype data were available for all mice, but the number of mice phenotyped for each organismal phenotype varied. Covariates such as sex, body weight and group size were considered and phenotypes normalised using a covariate-aware Box-Cox transformation. The residuals of a linear model including the covariates as fixed effects were used in all analyses of CFW data.

### Specific information about the blood and serum phenotypes collected in HS and CFW mice

In HS mice, blood was collected, first, in naive mice and analysed by FACS (“Imm.” phenotypes); then just before and at regular intervals after intraperitoneal glucose injection, and analysed with a glucose analyser (“Glucose” phenotypes); and finally after cardiac puncture for analysed using a medical-grade haematology analyser (“Haem” phenotypes) and an automated clinical chemistry analyser (“Biochem” phenotypes).

In CFW mice, blood was collected after cardiac puncture, for FACS, haematology and biochemistry analysis.

### Experimental design and confounding

Confounding between DGE, IGE and cage effects (in HS mice only): Because the average relatedness of co-housed mice is greater than the average relatedness of non co-housed mice in the HS dataset, DGE and IGE are partially confounded. To the extent that the cage environment affects the phenotypes studied here, they are also partially confounded with cage effects. However, we demonstrated both before^14^ and again in this study that accounting for DGE, IGE and cage effects jointly in our models of phenotypic variation yields unbiased estimates of DGE, IGE and IGE-DGE univariate and bivariate genetic correlations.

Confounding between DGE and parental effects (in HS and CFW mice): DGE could be partially confounded with parental effects, to the extent that the pre-weaning parental environment affects the adult phenotypes studied here.

Confounding between IGE and parental effects (in HS mice only): Because the average relatedness of co-housed mice is greater than the average relatedness of non co-housed mice in the HS dataset, IGE are partially confounded with parental effects, to the extent that the pre-weaning parental environment affects the adult phenotypes studied here. We checked that the IGE-DGE correlations analysed in this study were largely unaffected by this confounding (Supplementary Fig. 5).

### Simulations to validate our new implementation of bivariate IGE-DGE models (Supplementary Table 3 and Supplementary Fig. 1)

For each dataset, we simulated 1,000 pairs of “mock” phenotypes using the bivariate version of model (1), the real genetic relatedness matrix and cage assignments, and parameter values for the phenotypic covariance (3) that mimicked a favourable scenario with relatively large DGE and IGE, and intermediate correlation values (Supplementary Table 3). To do so, we used the mvn() function from the R MASS package.

We analysed each pair of simulated phenotypes the same way we analysed the real phenotypes and, for each parameter, examined the difference between the simulated and the estimated value of the parameter (Supplementary Fig. 1).

### Null simulations to validate *p*-values for 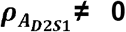 (Supplementary Table 4 and Supplementary Fig. 2)

We selected five phenotypic pairs from CFW mice with increasing p-values: “Haem.MPV”-“Haem.Large_PLT” (p-value of 1·10^-4^), “Bioch.LDL_BWcorr”-”Haem.Large_PLT” (p-value of 0.001), “Hypoxia.TV_SHR_BWcorr”-”Neuro.Ki67_BWcorr” (p-value of 0.01), “Haem.MPV”-“Bioch.Iron_BWcorr” (p-value of 0.1), and “Bioch.CreatinineEnzymatic_BWcorr”-“Bioch.ALP_BWcorr” (p-value of 0.99). The smallest p-value, 10^-4^, is close to the FDR-adjusted significance threshold (p ≤ 2.4·10^-4^). We first fitted the null model with 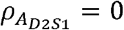 and generated null simulations, as described in the previous section, setting the parameter values to the parameter values estimates obtained from these null models. We simulated 10,000 pairs for the phenotype pair with the 1·10^-^4 *p*-value and 1,000 pairs for the other four phenotype pairs.

We analysed each pair of simulated phenotypes the same way we analysed the real phenotypes and derived LRT statistics and p-values for the hypothesis that 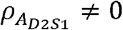.

To evaluate our p-value, we compared in a quantile-quantile plot the p-values from the null simulations to a uniform distribution of values between 0 and 1, which is the expected distribution of p-values if 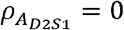 (Supplementary Fig. 2).

### Simulations to validate sex-specific bivariate IGE-DGE models (Supplementary Table 3 and Supplementary Fig. 3)

For each dataset, we simulated 1,000 “mock” phenotypes using a similar strategy to the one described above. We assigned missing values to males to simulate the female phenotype (*y*_1_), missing values to females for the male phenotype (*y*_2_) and finally concatenated them in a single phenotype. We used the same favourable scenario with large DGE and IGE, and intermediate correlation values (Supplementary Table 3).

We analysed each pair of simulated phenotypes the same way we analysed the real phenotypes and, for each parameter, examined the difference between the simulated and the estimated value of the parameter (Supplementary Fig. 3).

### Null simulations to validate *p*-values for 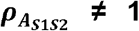 (Supplementary Table 5 and Supplementary Fig. 4)

We selected the phenotype from CFW mice with the smallest p-value: “Haem.EOS_percent” (p of 0.004). We first fitted the null model with 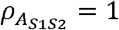 and generated null simulations, as described in the previous section, setting the parameter values to the parameter values estimates obtained from the null model of the real phenotype.

We analysed each phenotype the same way we analysed the real phenotypes and derived LRT statistics and p-values for the hypothesis that 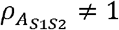.

To evaluate our p-value, we compared in a quantile-quantile plot the p-values from the null simulations to a uniform distribution of values between 0 and 1, which is the expected distribution of p-values if 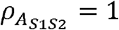 (Supplementary Fig. 4).

### Computational performance

The analysis of the phenotype pairs from the HS mice dataset took between 12 minutes and 19 hours and 25 minutes, using 1.2 to 4.9 GB of RAM. For CFW mice, runtime ranged from 14 minutes to 3 hours and 8 minutes, with RAM usage between 1.3 and 3.0 GB.

## Data availability

All data are available from published datasets. Details are provided on the mouse data used in this study in the Methods section. Data from Santostefano et al.^9^ were also re-analysed.

## Code availability

The computational pipeline used in this study is available from https://github.com/Baud-lab/CoreQuantGen.

## Acknowledgements

This work was supported by PID2021-122651NA-I00 and PRE2021-097413 contracts funded by MCIN/AEI/10.13039/501100011033 and FSE+ to AB and HT. We acknowledge support of the Spanish Ministry of Science and Innovation through the Centro de Excelencia Severo Ochoa (CEX2020-001049-S, MCIN/AEI /10.13039/501100011033), the Generalitat de Catalunya through the CERCA programme and to the EMBL partnership. FPC was funded by the Free State of Bavaria’s Hightech Agenda through the Institute of AI for Health (AIH).

## Author contributions

AB designed the study, with support from FPC. HT, FPC and AB developed the code. HT performed the analyses. HT, FPC and AB wrote the manuscript.

