## Supplementary tables and figures for "Phenome-wide genetic framework to identify mechanisms of social effects"

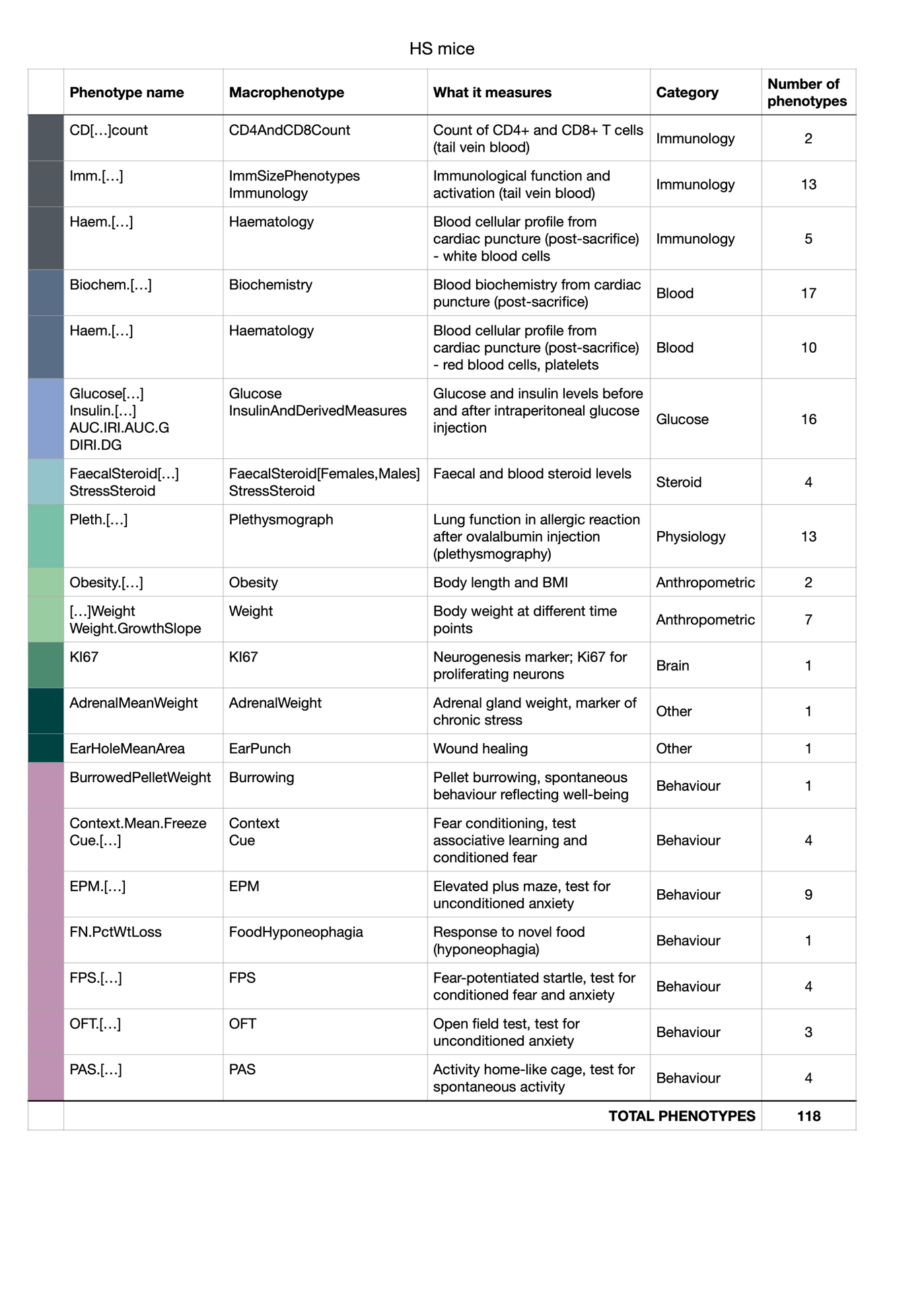

###### **Supplementary Table 1. Phenotypes measured in HS mice**

Comprehensive overview of 118 phenotypes measured in outbred HS mice. Phenotypes are organized and coloured by macro-phenotype spanning different phenotypic categories, capturing a variety of behavioural and non-behavioural traits. Each phenotype represents a measurement obtained from a test (timeline of test execution is shown in Fig. 2).

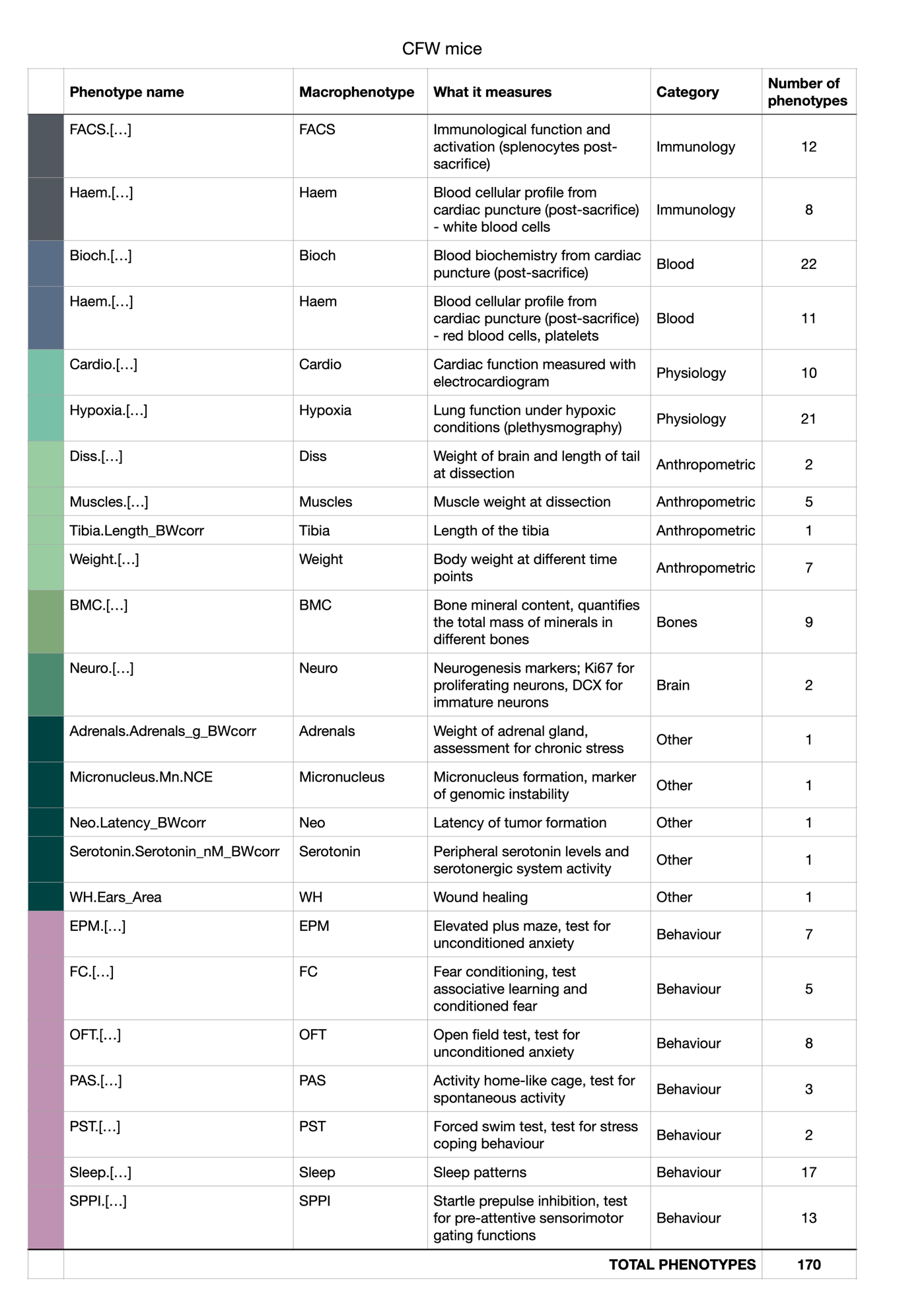

###### **Supplementary Table 2. Phenotypes measured in CFW mice**

Comprehensive overview of 170 phenotypes measured in outbred CFW mice. Phenotypes are organized and coloured by macro-phenotype spanning different phenotypic categories, capturing a variety of behavioural and non-behavioural traits. Each phenotype represents a measurement obtained from a test (timeline of test execution is shown in Fig. 2).

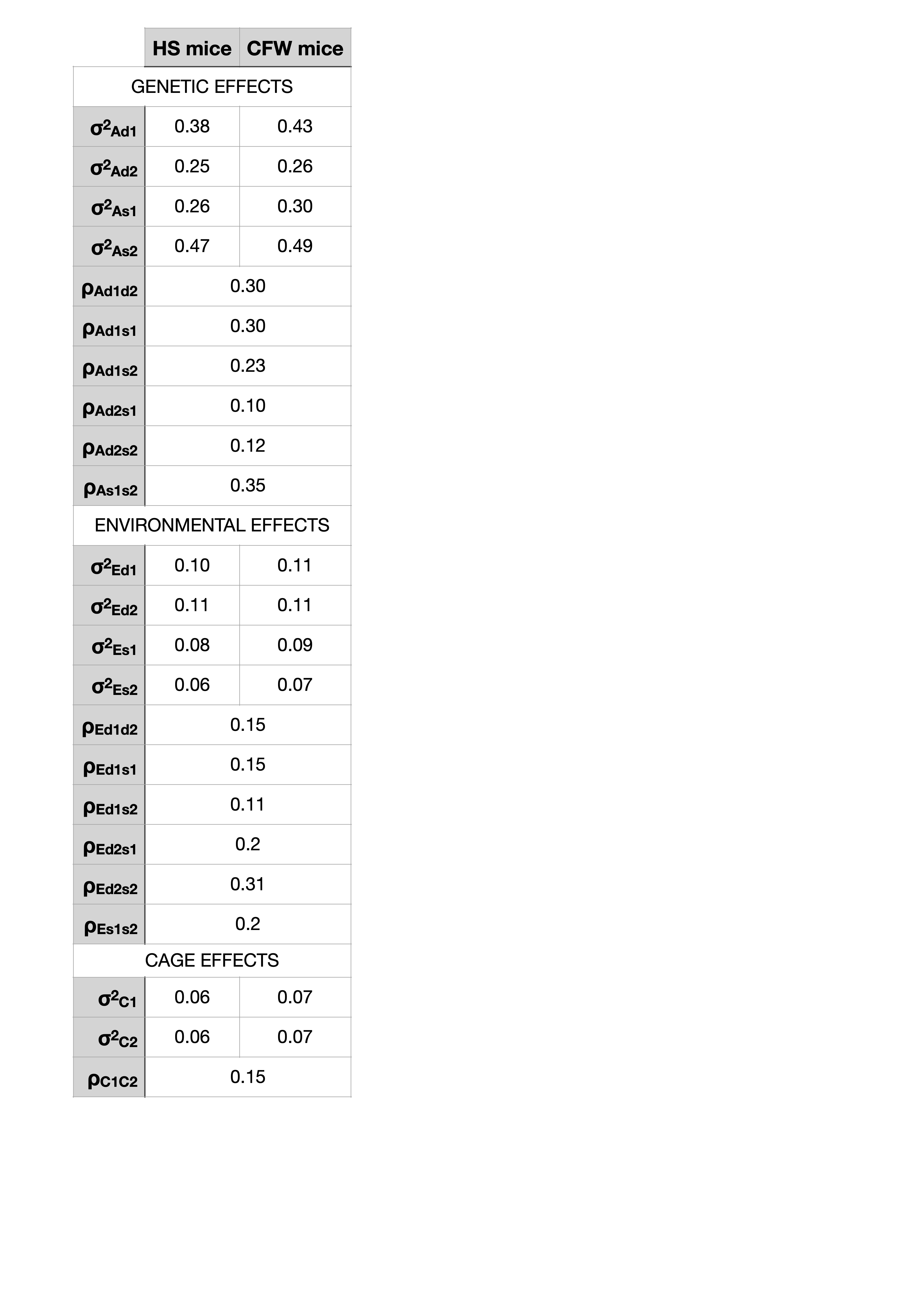

###### **Supplementary Table 3. Values set in simulation to validate the implementation of bivariate IGE-DGE models and sex-specific bivariate IGE-DGE models.**

For each parameter we report the value set in simulations in HS and CFW mice datasets. The same values were used to simulate pairs of “mock” phenotypes (Supplementary Fig. 1) and “mock” female/male phenotypes (Supplementary Fig. 3).

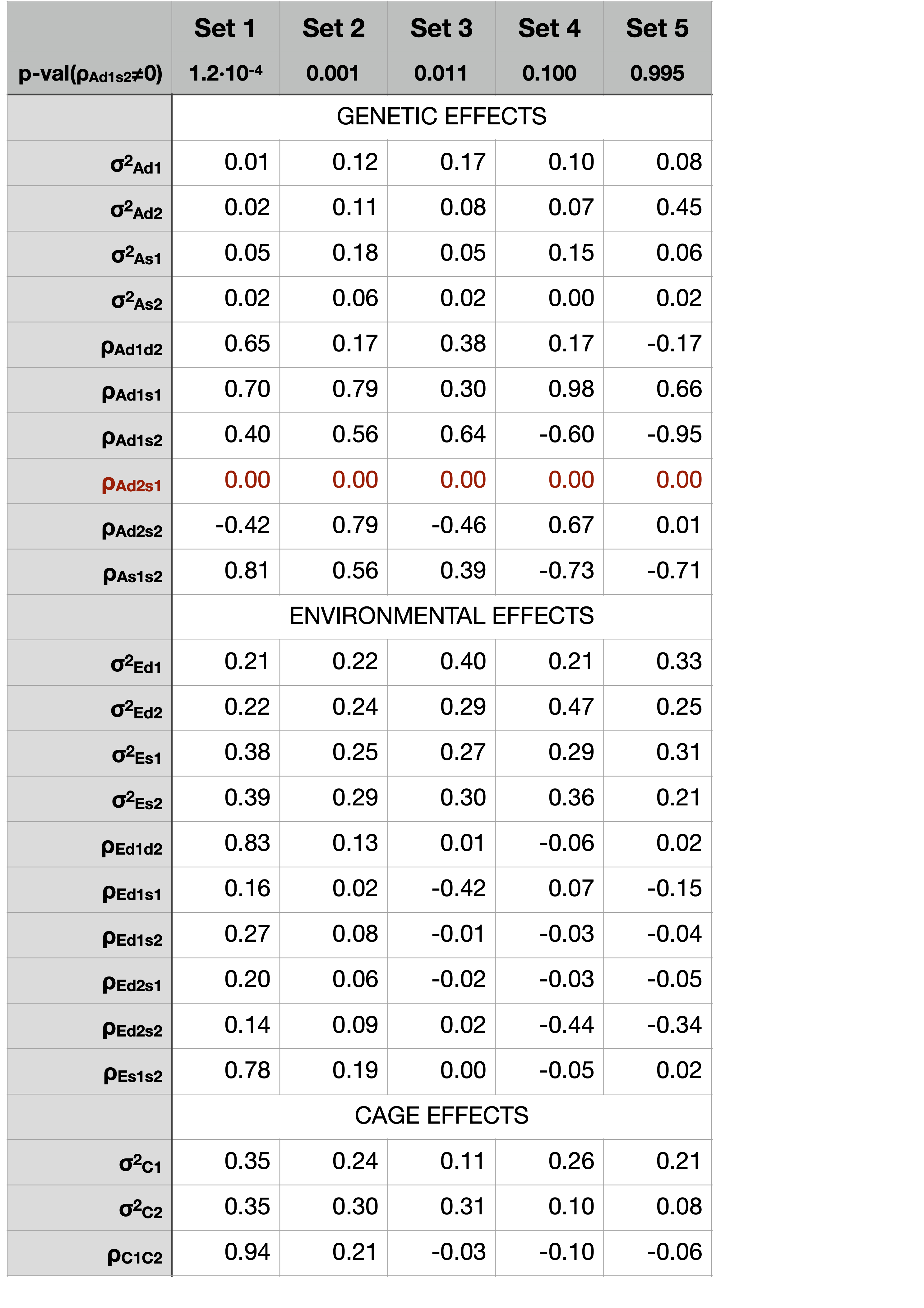

###### **Supplementary Table 4. Parameter values used in null simulations to validate p-values for** $\boldsymbol{\rho}_{\boldsymbol{A}_{\boldsymbol{D}\boldsymbol{2}\boldsymbol{S}\boldsymbol{1}}}$**≠ 0.**

For each parameter we report the values set to simulate pairs of “mock” phenotypes under the null hypothesis of $\rho_{A_{D2S1}}=0$. The values were derived from the null models of five phenotypic pairs with increasing p-values, selected from CFW mice: “Haem.MPV”- “Haem.Large_PLT” (set 1), “Bioch.LDL_BWcorr”-“Haem.Large_PLT” (set 2), “Hypoxia.TV_SHR_BWcorr”-“Neuro.Ki67_BWcorr” (set 3), “Haem.MPV”-“Bioch.Iron_BWcorr” (set 4), and “Bioch.CreatinineEnzymatic_BWcorr”-“Bioch.ALP_BWcorr” (set 5).

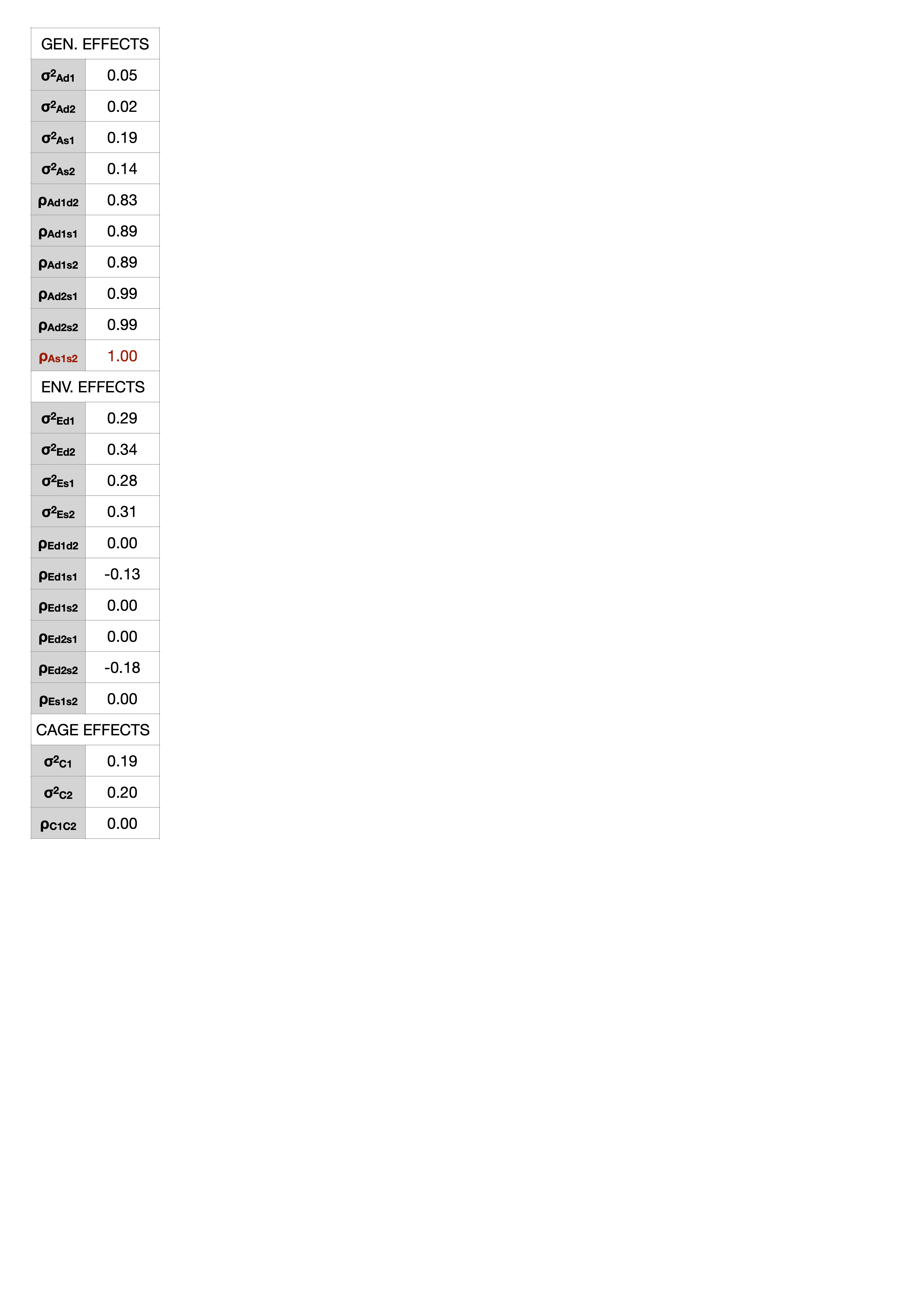

###### **Supplementary Table 5. Parameter values used in null simulations to validate p-values for** $\boldsymbol{\rho}_{\boldsymbol{A}_{\boldsymbol{S}\boldsymbol{1}\boldsymbol{S}\boldsymbol{2}}}$**≠ 1 (sex-specific bivariate IGE-DGE model).**

For each parameter we report the value set to simulate “mock” female/male phenotypes under the null hypothesis of $\rho_{A_{S1S2}}=1$. The values were derived from the null model of a selected phenotype from CFW mice (“Haem.EOS_percent”, p-value of 0.004).

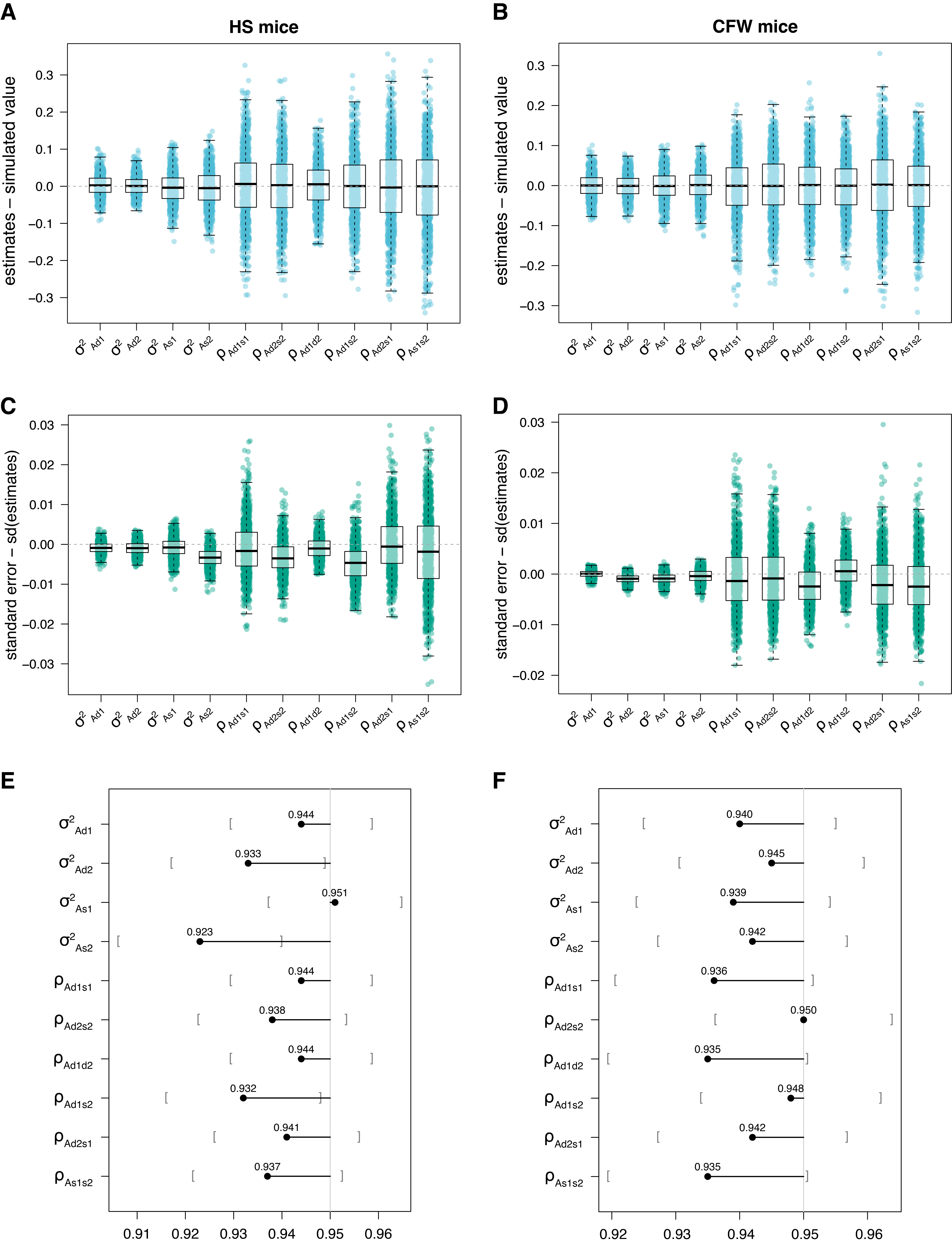

###### **Supplementary Figure 1. Validation of genetic parameter estimates using simulated data.**

###### (A,C,E) HS mice. (B,D,F) CFW mice. (A) and (B) For each genetic parameter shown on the x-axis, the y-axis shows the differences between estimated and simulated value (Supplementary Table 3) across 1,000 simulated phenotype pairs. Each dot represents one phenotypic pair, and boxplots summarize their distributions. Subscripts indicate phenotype indexing: index 1 denotes the phenotype of interest measured in focal individuals, whereas index 2 denotes a measured phenotype whose quality as a proxy phenotype is being evaluated. (C) and (B) For each genetic parameter shown on the x-axis, the y-axis shows the differences between estimated and empirical standard error across 1,000 simulated phenotype pairs. Each dot represents one phenotypic pair, and boxplots summarize their distributions. (E) and (F) For each genetic parameter a lollipop bar shows the probability that the simulated parameter value was within 95% confidence interval, calculated as the value of the estimate +/- 1.96 x the standard error. Grey square brackets show Monte Carlo 95% confidence interval for the probability estimate.

**Note for interpretation:**

Using simulations based on the real genotypes and cage assignments of HS and CFW mice and under a scenario of relatively strong genetic effects (IGE explaining 26% and 30% of phenotypic variance for phenotype 1 and DGE explaining 25% and 26% for phenotype 2 in HS and CFW mice respectively), we confirmed that all the genetic parameters were unbiased (mean difference between simulated and observed value lower than 1%, Supplementary Fig. 1A and 1B) and that the estimated standard errors were very close to the empirical standard error (i.e. standard deviation of the estimates) albeit slightly smaller (Supplementary Fig. 1C and 1D). As a result, the simulated parameter value was within the 95% confidence interval, calculated as the value of the estimate +/- 1.96 x the standard error, in 92 to 95% and in 93 to 95% of cases in HS mice and in CFW mice respectively (Supplementary Fig. 1E and 1F).

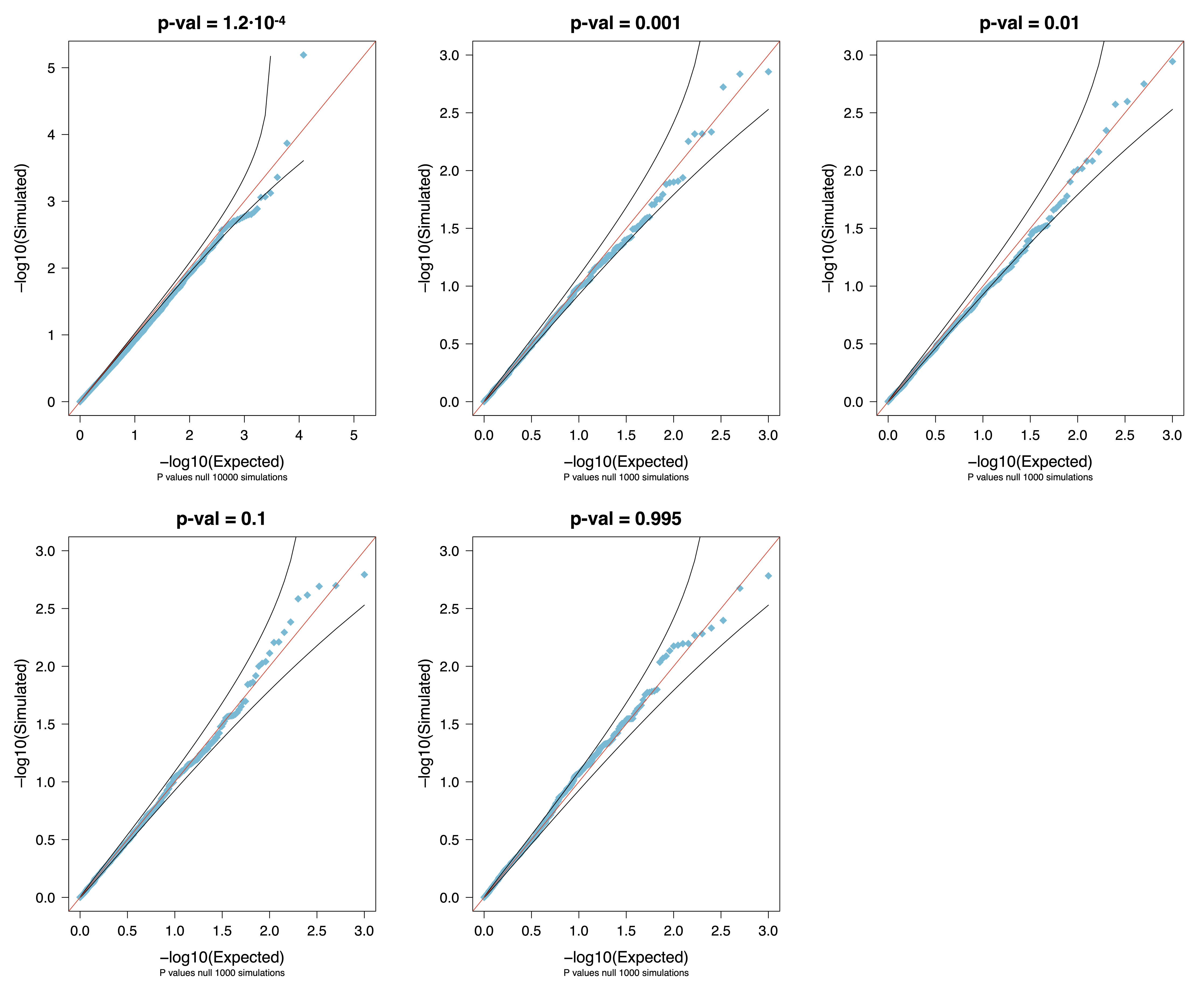

### Supplementary Figure 2. Calibration of *p*-values for $\boldsymbol{\rho}_{\boldsymbol{A}_{\boldsymbol{D}\mathbf{2}\boldsymbol{S}\mathbf{1}}}$≠ 0 using null simulations.

Each panel shows the Q-Q plot of simulated p-values (blue dots) compared to the expected uniform distribution (red diagonal line). Black lines around the diagonal represent 95% confidence intervals under the null hypothesis. Deviations from the diagonal would indicate miscalibration. The plots correspond to simulations based on phenotype pairs from CFW mice, with observed p-values ranging from 1.2·10⁻⁴ to 0.995 (Supplementary Table 4).

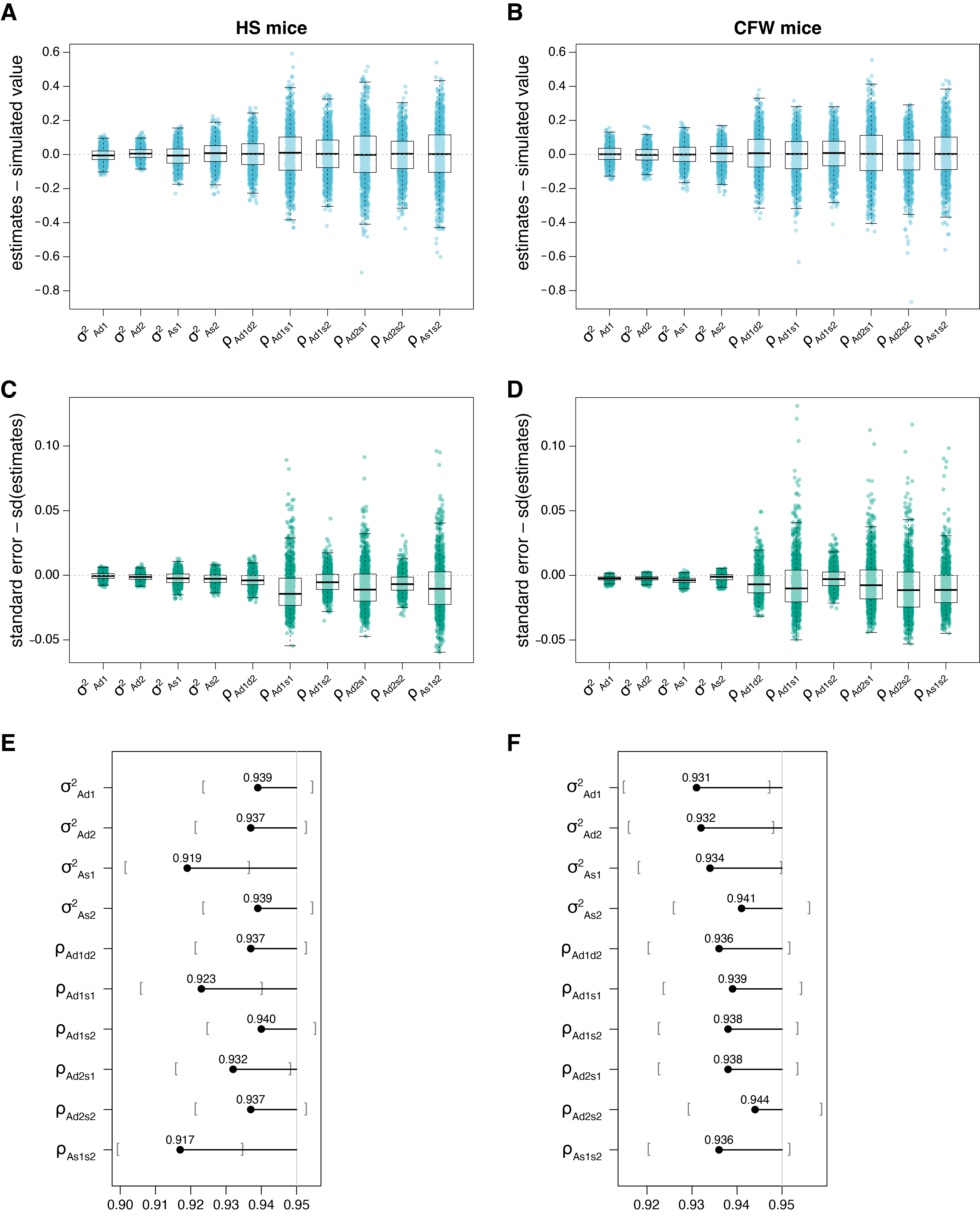

### Supplementary Figure 3. Validation of genetic parameter estimates using simulations (sex-specific bivariate IGE-DGE model).

###### (A,C,E) HS mice. (B,D,F) CFW mice. (A) and (B) For each genetic parameter shown on the x-axis, the y-axis shows the differences between estimated and simulated value (Supplementary Table 3) across 1,000 simulated phenotypes. Each dot represents one phenotype, and boxplots summarize their distributions. Subscripts indicate phenotype indexing: index 1 denotes a female phenotype (males had missing values), whereas index 2 denotes a male phenotype (females had missing values). (C) and (B) For each genetic parameter shown on the x-axis, the y-axis shows the differences between estimated and empirical standard error across 1,000 simulated phenotypes. Each dot represents one phenotypes, and boxplots summarize their distributions. (E) and (F) For each genetic parameter a lollipop bar shows the probability that the simulated parameter value was within 95% confidence interval, calculated as the value of the estimate +/- 1.96 x the standard error. Grey square brackets show Monte Carlo 95% confidence interval for the probability estimate.

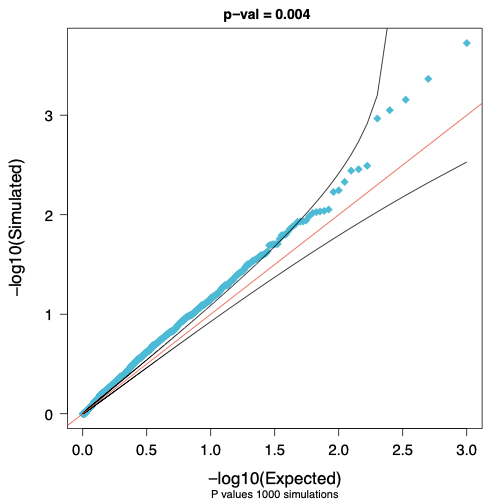

### Supplementary Figure 4. Calibration of *p*-values for $\boldsymbol{\rho}_{\boldsymbol{A}_{\boldsymbol{S}\mathbf{1}\boldsymbol{S}\mathbf{2}}}$≠ 1 using null simulations (sex-specific bivariate IGE-DGE model).

Q-Q plot of simulated p-values (blue dots) compared to the expected uniform distribution (red diagonal line). Black lines around the diagonal represent 95% confidence intervals under the null hypothesis. Deviations from the diagonal would indicate miscalibration. The plots correspond to simulations based on a selected phenotype from CFW mice (Supplementary Table 5).

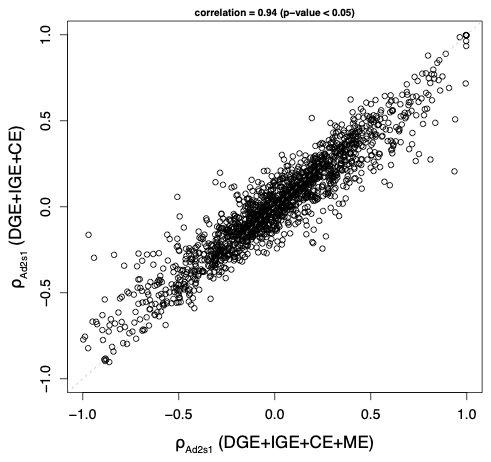

**Supplementary Figure 5. IGE-DGE correlations analysed in this study (**$\boldsymbol{\rho}_{\boldsymbol{A}_{\boldsymbol{D}\mathbf{2}\boldsymbol{S}\mathbf{1}}}$**) are largely unaffected maternal effects.**

Each dot represents a pair of phenotypes in HS mice. Dots distribute along the x-axis depending on the value of the IGE-DGE correlation ($\rho_{A_{D2S1}}$) estimated under a model including maternal effects and along the y-axis depending on the value of the same correlation estimated under a model excluding maternal effects (these correspond to the values presented in the main text, Fig. 4 and Fig. 5). Estimates are significantly correlated with a value of 0.94 (*p*-value < 0.05).
